# CD15 and CD15s expression is associated with G1 phase of the cell cycle in glioma cell lines

**DOI:** 10.1101/2020.11.17.387019

**Authors:** Samah A. Jassam, Zaynah Maherally, Paraskevi Chairta, Geoffrey J. Pilkington, Helen L. Fillmore

## Abstract

Overexpression of the tetrasaccharide carbohydrate epitopes, CD15 and CD15s are associated with non-central nervous system malignancies. While CD15 and CD15s expression is rare in gliomas, recent reports suggest that CD15 may serve as a marker for brain tumour ‘stem-like’ cells. The aim of this study was to determine if this apparent discrepancy may, in part, be explained by temporal expression of CD15 and CD15s at different phases of the cell cycle. We used flow cytometry, immunocytochemistry and a fluorescence cell cycle indicator (FUCCI) system to examine expression in glioblastoma (GBM) cells (UP-007 and SNB-19) and non-neoplastic astrocytes (SC-1800) synchronised via serum starvation, Hydroxyurea and Nocodazole, respectively. CD15 and CD15s expression was significantly increased in glioma cells synchronised to G1 phase compared with non-synchronised cells (p<0.001). This was supported by qualitative results obtained with the (FUCCI) system. Few studies have considered the possibility of cell-cycle dependent CD15 and CD15s expression which may explain the inconsistencies reported in the literature in terms of expression in ‘glioma stem-like cells’ where cells are more likely in S phase where CD15 and CD15s expression would be low.

## Introduction

The expression of cell membrane glycoconjugates is reported to be highly influenced by cell differentiation status [1, 2]. Some cancer cells express particular glycoconjugates during the transformation process and the overexpression of some of these conjugates has been directly correlated with malignancy [3]. CD15 (Lewis x) (Galβ1,4[Fucα1,3] GlcNAc), a trisaccharide molecule also known as a Stage-Specific Embryonic Antigen-1 (SSEA-1) [4], was described for the first time in a patient with red blood cell incompatibility and assigned after the patient’s family name “Lewis” [5]. Sialyl CD15 (CD15s) (α-Neup5Ac-(2, 3)-β-Galp-(1, 4)[α-Fucp-(1,3)]-GlcpNAc-R) is a tetrasaccharide epitope that was first detected in a ganglioside fraction from human kidney [6] and in human milk [7]. CD15s has been shown to play a key role in cell-cell recognition in sperm-egg fertilisation and in extravasation of leukocytes [9]. Both CD15 and CD15s are expressed in a variety of non-neoplastic cells such as granulocytes, monocytes [10], epithelial cells [11] and neurons [12, 13]. A considerable number of studies have shown a strong correlation between overexpression of CD15, CD15s and malignancy in different cancers such as; Hodgkin’s lymphoma [14], breast [15, 16], lung [17], renal carcinoma [18], colon [19], head and neck squamous carcinoma [20] as well as brain metastases from lung [21, 17] and breast [16]. Although CD15 was previously reported to be rarely expressed on anaplastic glioma and glioblastoma [12, 22], it has also been suggested that CD15 may serve as a marker for brain cancer stem-like cells in human gliomas [23, 24, 25, 26]. Others have reported that CD15 expression is not a marker for glioma ‘stem-like’ cells [27]. The aim of this study was to gain a better understanding of CD15 and CD15s expression in GBM. Herein we demonstrate that the expression of CD15 and CD15s is cell cycle dependent in the glioma cell lines tested.

## Results

### Expression of CD15 and CD15s in non-synchronised brain tumour cells

Expression of CD15 and CD15s were characterised in non-synchronised cell cultures as per the flow chart described in Figure 1. Under normal (non-synchronised) culture conditions, immunocytochemistry and confocal results indicate an absence of CD15 expression in non-neoplastic astrocytes (SC-1800), while in glioblastoma cells (SNB-19 and UP-007), expression of CD15 was noted on the cell surface of individual cells (Figure 2a). Semi-quantitative analysis revealed a 5 fold higher number in CD15 fluorescence intensity in UP-007 cells and a 4 fold higher in SNB-19 cells compared to SC-1800 and isotype control (Figure 2c, p< 0.05). Similarly, CD15s was not detected on the surface of the non-neoplastic astrocyte cell culture, SC-1800 whereas a faint expression was detected on some glioma-derived cells (Figure 2b). As with CD15, CD15s was localised to the outer edges of the cell in monolayer cultures. Semi-quantitative analysis demonstrated a 2.9 fold increase in CD15s expression in UP-007 and a 5.1 fold increase in SNB-19 cells compared to the isotype controls (Figure 2d, p<0.05).

**Figure 1:**
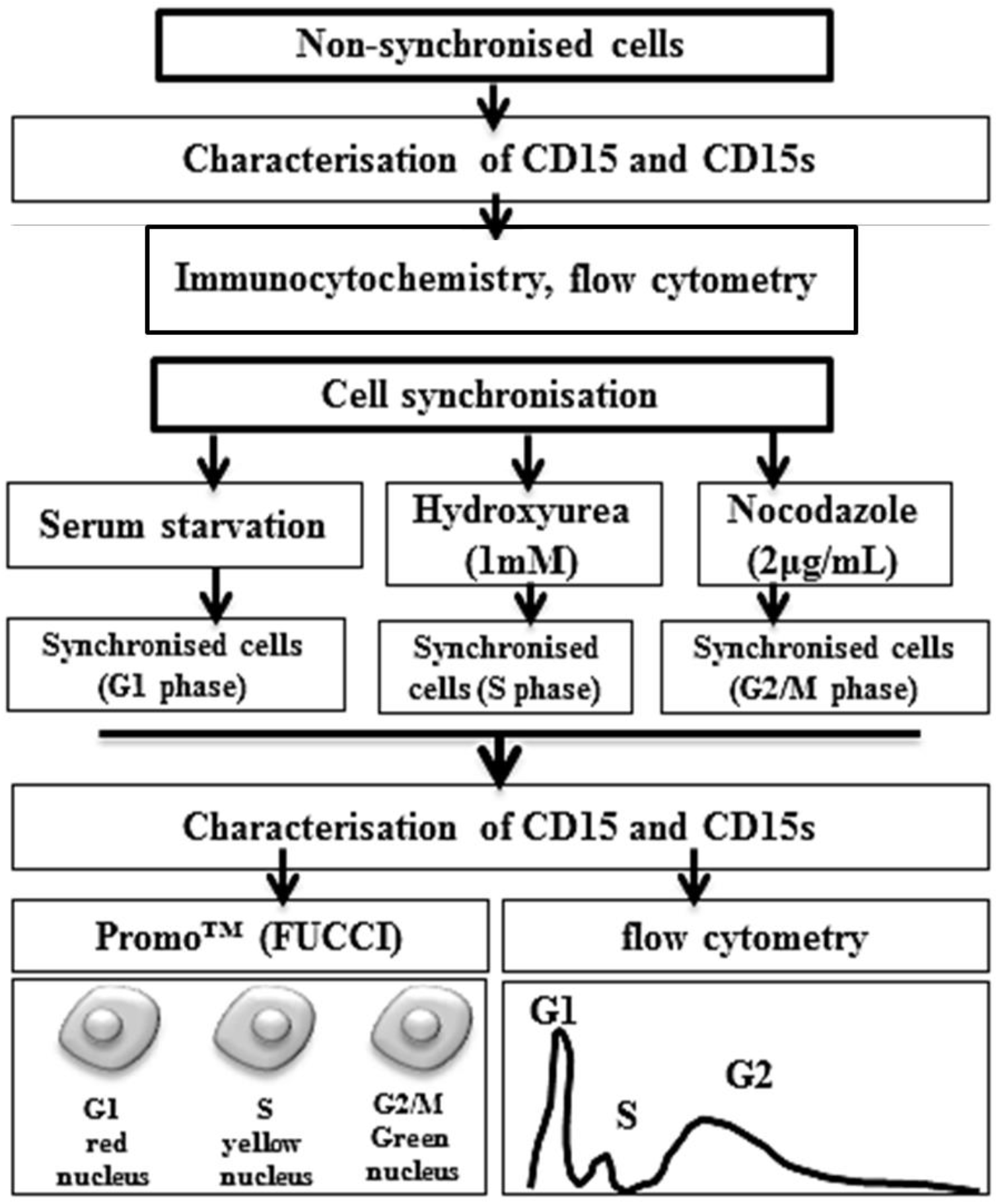
Schematic diagram showing the methodology for characterisation of CD15 and CD15 at different stages of cell cycle. CD15 and CD15s expression was characterised in non-synchronised cell cultures in glioma cell lines (UP-007 and SNB-19) and an astrocyte cell line (SC-1800) using immunocytochemistry and flow cytometry analysis. Cell cultures were then synchronised at G1 phase by serum starvation, at S phase via Hydroxyurea (1mM) and at G2/M phase via Nocodazole followed by characterisation of CD15 and CD15s in the synchronised cells using ICC/Promo™ (FUCCI) and flow cytometry analysis.

**Figure 2:**
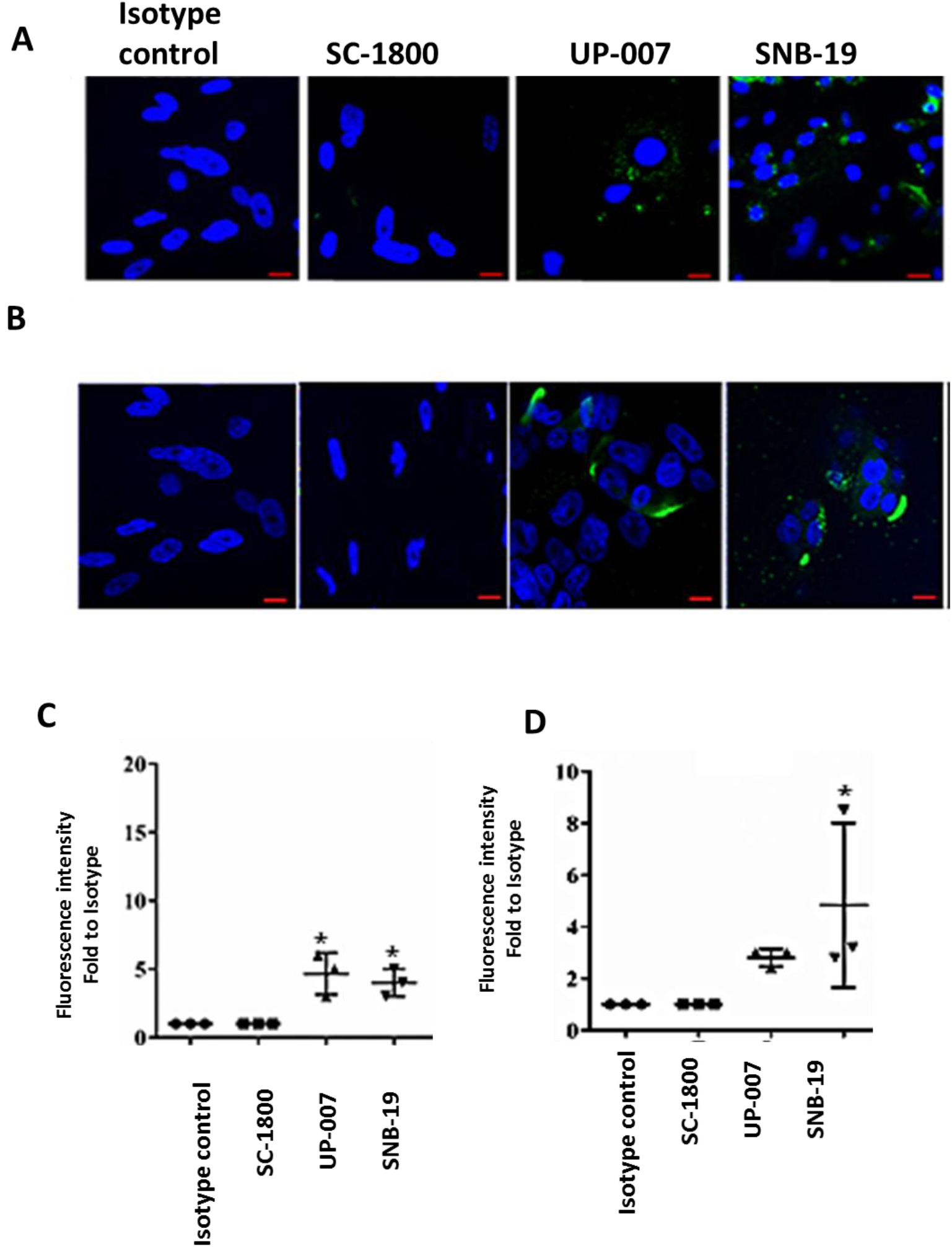
Expression and localisation of CD15 and CD15s in non-synchronised cells. Immunocytochemistry images showing no CD15 **(a**) and CD15s **(b)** expression (green) in non-neoplastic astrocytes (SC-1800) but positivity in glioma cells (UP-007 and SNB-19). Hoechst blue was used as a nuclear counterstain. Scale bar =20μm. Semi-quantification analysis of CD15 **(c)** and CD15s **(d)** from confocal microscopy images using Zeiss Zen image software.

### Expression of CD15 at different cell cycle stages

CD15 and CD15s expression levels were assessed in the two glioma cell lines (UP-007 and SNB-19) as well as in the astrocyte cells (SC-1800), synchronised at ether G1, S or G2/M (Figure 1). Prior to this analysis, of the relationship between cell cycle phases and CD15 and CD15s expression, the efficiency of synchronisation was determined in each in cell line using DNA analysis via flow cytometry. Cells were synchronised to G1, S and G2/M phases of the cell cycle using serum depletion, hydroxyurea and Nocodazole respectively and compared with control non-treated cells (Supplemental Figure 1). In non-synchronised conditions, most of the SC-1800 cells were in G2/M phase while in glioma cell lines, most were in G1 phase as determined by flow cytometry analysis of propidium iodide stained cells (Supplemental Figure 1a). In terms of CD15 expression in non-synchronised cells, flow cytometry analysis indicated that only 3% of SC-1800 expressed CD15 and this was not significantly different from isotype controls (Supplemental Figure 2a).

In synchronised cells, flow cytometry analysis revealed that CD15 was expressed at low levels in SC-1800 with no observable difference in expression in G1 (1.8%), S (1.18%) or G2/M (1%) while in GBM cells, CD15 expression was significantly higher in cells arrested at G1 phase (14.7% and 21.4% in UP-007 (p<0.05) and SNB-19 (p<0.01) respectively compared to non-synchronised counterparts) (Figure 2 a and c). In addition, in each of the GBM cell lines there was significantly higher CD15 expression in G1 compared to S (p<0.01) and G2/M (p<0.01). These data were supported by qualitative results obtained from transfected cells using the Premo™ (FUCCI) cell cycle sensor to differentiate specific cell cycle stages, visualised using a LSM 510 Axioskop2 confocal microscope. G1, S and G2/M phases were detected by red, yellow and green nuclei respectively (Figure 3a and b). In these non-synchronised studies, CD15 cell surface expression (green) was mostly co-localised with red nuclei indicative of G1 (Figure 3c) supporting that higher levels of CD15 expression can be detected in cells at G1 phase of the cell cycle.

**Figure 3:**
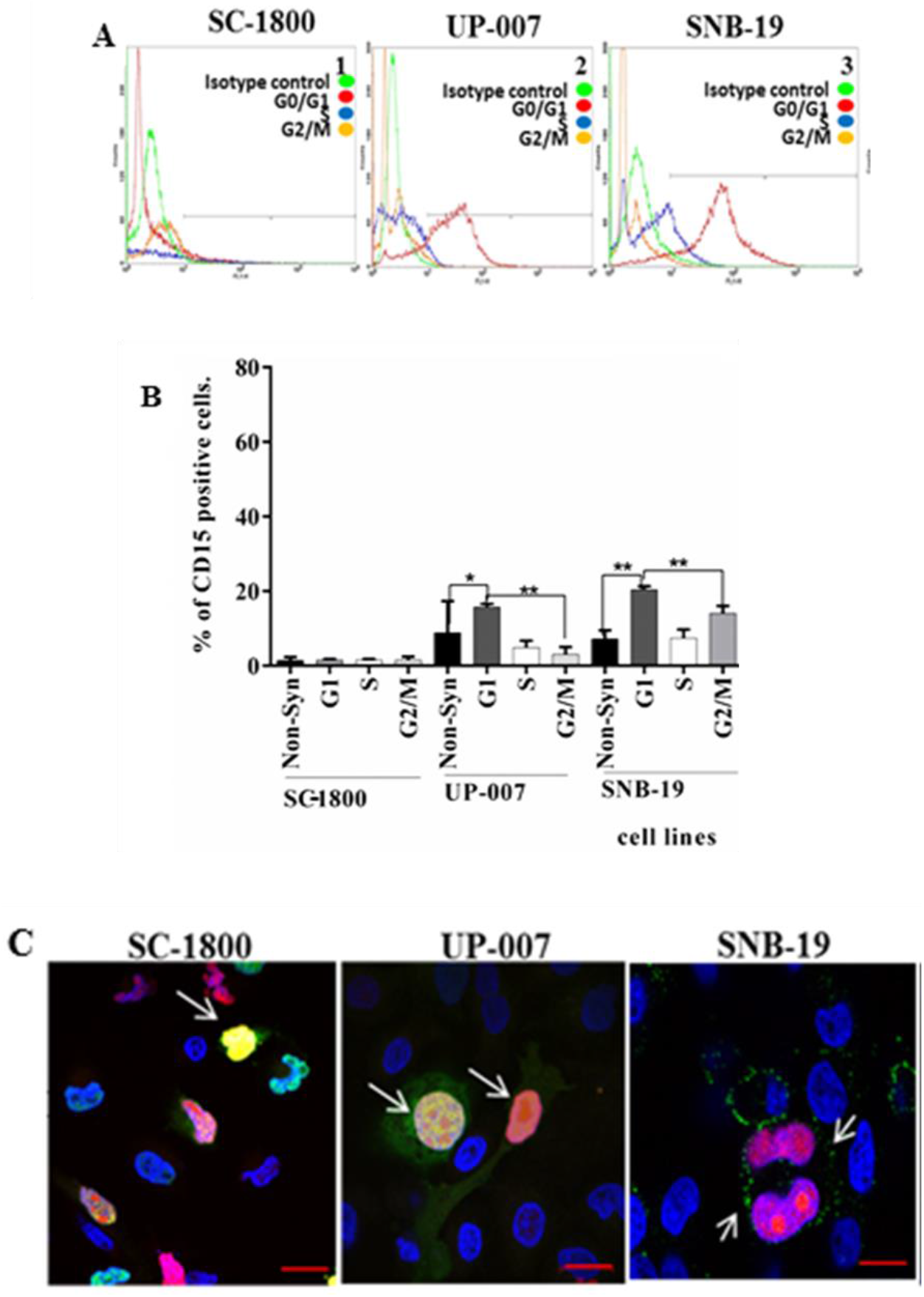
Expression and localisation of CD15 during different stages of the cell cycle. **(a)** Flow cytometric histograms representing CD15 expression in synchronised cells SC-1800 (1), UP-007 (2), SNB-19 (3). CD15 positive expression is represented by a shift of the graph to the right. **(b)** Flow cytometric analysis showing the highest level of CD15 expression was detected in G0/G1 synchronised cells followed by non-synchronised cells and G2/M phase compared to IgM isotype control. Cells arrested at S phase showed least CD15 expression level. **(c)** Immunocytochemistry results of CD15 extracellular expression (green). Cells were treated with FUCCI™ system to differentiate specific cell cycle stages depending on the colour of the nuclei (red: G1; yellow: S and green: G2/M phase). Results show CD15 expression was detected at low levels on SC-1800 cell with red nuclei (G1 phase). In GBM cells (UP-007 & SNB-19), CD15 was expressed and well distributed on cell surfaces with red or yellowish-red nuclei (arrow) indicating G1 phase *(P≤0.05) and ** (P<0.01). Scale bar is 20μm. Results are representative of three independent experiments carried out in triplicate (n=3).

### Expression of CD15s at different cell cycle stages

CD15s expression also shows cell-cycle dependency. In non-synchronised cells, 2% of SC-1800 cells, 8% of SNB-19 cells and 16% of UP-007 cells were CD15s positive (Supplemental Figure 2b). CD15s was expressed at a lower level in SNB-19 (9.5%) and UP-007 (13.4%) compared to isotype control (p<0.05 and p<0.01). As with CD15, CD15s expression in the astrocyte cell line was low and did not significantly change with synchronisation. In GBM cells, CD15s expression was significantly higher in cells arrested at G1 phase: 39% and 48% in UP-007 (p<0.001) and SNB-19 (p<0.001) respectively compared to non-synchronised counterparts (Figure 4a and b). In addition, in each of the GBM cell lines there was significantly higher CD15s expression in G1 compared to S (p<0.001) and G2/M (p<0.001). The increase in CD15s expression in glioma cell lines synchronised to G0/G1 was dramatically higher than CD15 expression. In non-synchronised co-localisation studies, CD15s expression (green) was also co-localised with red nuclei indicative of G1 (Figure 4c). These results suggest that expression of CD15s correlates with G1 phase.

**Figure 4:**
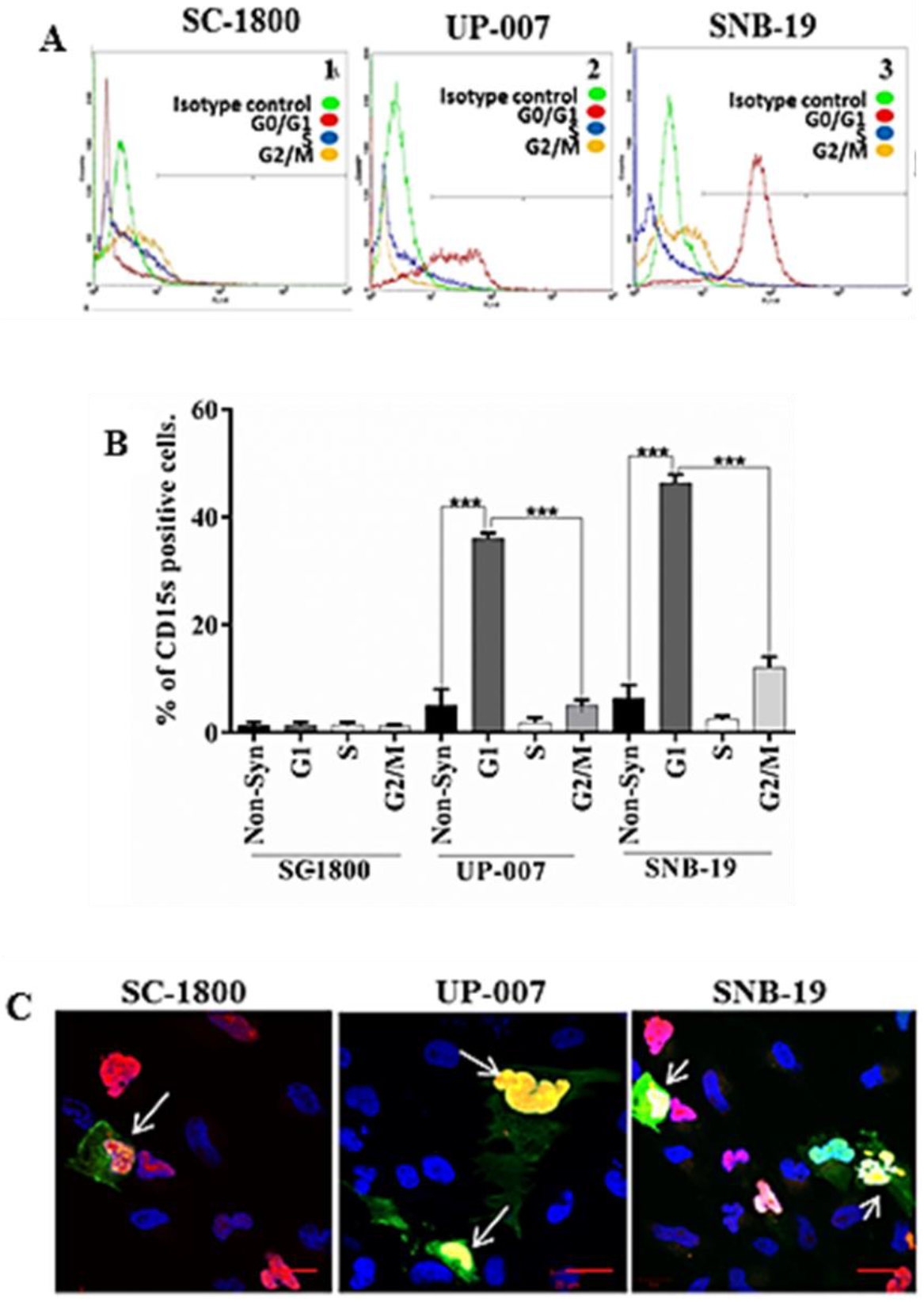
Expression and localisation of CD15s during different stages of the cell cycle. **(a)** Flow cytometric histograms representing CD15s expression in synchronised cells: SC-1800 (1), UP-007 (2), SNB-19 (3) cells. CD15s positive expression is represented by a shift to the right. **(b)** Flow cytometric analysis show highest level of CD15s expression was detected in G0/G1 synchronised cells followed by non-synchronised cells and G2/M phase. Differences compared to the IgM isotype control. **(c)** Immunocytochemistry results of CD15s extracellular expression (green). Cells were treated with the FUCCI™ system to differentiate specific cell cycle stages depending on the colour of the nuclei (red: G1; yellow: S and green: G2/M phase). Results showed that CD15s expression was detected at low level on SC-1800 cells with red nuclei (G1 phase). In GBM cells (UP-007 & SNB-19), CD15s was expressed and well distributed on cell surfaces with red or yellowish-red nuclei (arrow) indicating G1 phase. Scale bar=20μm. Cells which arrested at S phase showed lowest CD15s expression levels *** (P<0.001). Results are representative of three independent experiments carried out in triplicate (n=3).

## Discussion

CD15 and CD15s are fucosylated polysaccharide epitopes and tumour-associated cell adhesion molecules. Overexpression of both epitopes has been correlated with malignancy of many non-CNS cancers [15, 14]. Early studies in CNS tumours reported the absence or low CD15 expression in GBM cells [12, 22]. However more recently, expression of CD15 in glioblastomas and specifically with tumour-initiating cells [23, 24, 28, 25, 26]. In contrast, Kenney-Herbert *et al* [27] reported that there were no phenotypic or genetic differences between CD15- and CD15+ GBM cells and CD15 expression was not enough to distinguish a discrete population of GBM cells. Mao *et al* [24] showed that in 20 GBM cases in which CD15 expression was investigated by immunohistochemistry, 12 cases were considered CD15 positive. In another study where CD15 expression was being investigated as a potential marker for rare extracranial metastases, 5/13 of GBM cases were considered CD15 positive [29]. Few studies have however, considered the possibility of cell-cycle dependent CD15 expression which may explain the inconsistencies reported in the literature. A recent study, however; showed elegantly, data to support the idea of an intrinsic glioma cancer stem cell plasticity which might help to explain the ‘inconsistencies’ [30].

CD15s expression has been shown to be variable in different non-synchronised GBM cell lines with positivity roughly from 4.8%-32.8% [31]. This agrees with our findings that show the variability in expression in non-synchronised GBM cell lines (UP-007 and SNB-19). In non-neoplastic astrocytes, CD15 and CD15s expression was low and this did not change when cell lines were synchronised to G1, S, or G2/M consistent with reports showing low CD15 expression in astrocytes [22].

In the GBM cell lines tested, the expression of CD15 and CD15s was significantly higher when cells are synchronised in G1 phase. These findings suggest that expression of both CD15 and CD15s in GBM cells correlate with G1 phase. Our overall findings suggest that in GBM cell lines, CD15 and CD15s expression is correlated with a specific stage of the cell cycle. This may help to explain the conflicting reports in the literature concerning CD15 expression in glioma cells and ‘glioma stem-like cells’ as differences could be due to majority of the cells being in the S phase where CD15 and CD15s expression would be low. Future studies are needed to address if this is the case. An additional interesting question is whether CD15 and CD15s play a role during cell cycle progression, particularly while cells are in G1. It has been suggested since the late 1980’s that cells in G1 were more susceptible to initiate differentiation in response to growth factors [32]. The concept of connecting the cell cycle phase and cell fate as well as the time spent in each phase has gained supportive experimental evidence over the years [reviewed in 33, 34]. Recently Singh [35] proposed a hypothetical model of the relationship between cell cycle, pluripotent cells and heterogeneity and that G1 could serve as a ‘differentiation induction point’ [35] and as proposed by Hardwick *et al* [33] the length of time ‘glioma stem-like cells’ are in G1 be targeted to enhance differentiation and slow tumour progression? How does the glioma ‘cancer stem cell niche’ contribute to cell cycle length? In an interesting proteomic study using a breast cancer cell line arrested in G1, bioinformatic tools and databases, Tenga and Lazar [36] reported that three major clusters of interacting networks emerged and included oxidative phosphorylation, DNA repair and signalling. These reports highlight the complexity associated with labelling a subset of cells. Instead of relying on ‘glioma stem-like cell’ markers the idea of using molecular regulatory components that act within a network may prove promising in terms of understanding and identification of therapeutic targets for GBM and ‘stem-like cells’. Two reports that examined CD15 as a potential glioma ‘stem-like’ cell also conducted CD15 IHC studies [23, 24]. The data presented herein on expression of CD15 being correlated with specific cell cycle phases are based on *in vitro* studies and future work should include *in vivo* experiments to determine if this phenotype is the same. Future work should also include a comprehensive temporal and spatial investigation of CD15 and CD15s in GBM and other brain tumour biopsy material. Interestingly, Qazi *et al* [25] demonstrated that the CD15 positive brain tumour stem-like cell population obtained from primary GBM can be enriched following chemo radiotherapy *in vitro* and this model may represent the phenotype seen in recurrent GBM. Exploring CD15 and CD15s expression, regulation and function in light of recent findings of CD15 as a potential glioma ‘stem-like’ marker with this work demonstrating the expression of these molecules are dependent on cell cycle phase, will provide additional valuable insight and hopefully lead to potential new therapeutic targets.

## Materials and methods

### Cell culture

This study included low passage adult human astrocytes (SC-1800) (passage 3-6) derived from the cerebral cortex (Caltag Medsystems, UK) and low and high passage glioblastoma multiforme, Grade IV (GBM) cells, UP-007 (passage 8-13) and SNB-19 (passage 38-42), respectively. UP-007 cells were established ‘*in house’* from biopsies received from surgical resections while SNB-19 was purchased from the DSMZ German Brain Tumour Bank, Germany. All cells were examined for mycoplasma contamination on a routine quality control basis, using MycoAlert™ kit (Lonza, Germany) and genetically authenticated using a STR-PCR fragments kit (Agilent Technologies, USA) as per our previous published technique [37]. SC-1800 cells were grown in astrocyte basal medium (ABM) supplemented with astrocyte single Quots™ (AGM-2) (Lonza, Germany) and 3% human serum (Sigma, UK). All other cell lines were grown in Dulbecco`s modified Eagle medium (DMEM) (Fisher, UK) supplemented with 2% human serum (Sigma, UK) and maintained in an humidified sterile incubator with 5% CO_2_ at 37°C.

### Antibodies

#### Primary antibodies

Mouse monoclonal anti-CD15 (MEM-158) IgM (Sigma, UK) and mouse monoclonal anti-sialyl CD15 (CD15s) (KM93) IgM (Millipore, UK) were used at the following dilutions: 1:100 and 1:50 for immunocytochemistry (ICC) and 1:10 and 1:10 for flow cytometry (FC) respectively.

#### Secondary antibodies

Fluorochrome-conjugated Alexa Fluor IgM 647nm (Invitrogen, USA) was used for ICC and flow cytometry at 1:500.

#### Isotype negative controls

Isotype negative control antibodies were used to confirm specificity of primary antibodies. Mouse monoclonal IgM Isotype antibodies (Invitrogen, USA) were used at the same concentrations of primary antibodies mentioned above.

### Synchronization of cell cultures at cell cycle specific stages

Expression of CD15 and CD15s were characterised in non-synchronised cell cultures in primary and secondary brain tumour cells as per the flow chart described in Figure 1. Cell cultures were synchronised at G1 phase by serum starvation, at S phase via Hydroxyurea (1mM) and at G2/M phase via Nocodazole (2μg/ml).

#### G1 phase

cells were arrested at G1 phase by serum deprivation. 1×10^6^ cells were seeded in T25 tissue culture flasks containing serum supplemented growth medium until 50% confluency was reached. Cells were then washed with pre-warmed sterile Hank’s buffered salt solution (Fisher, UK) followed by addition of growth medium supplemented with 1% human serum followed by overnight incubation. Cells were then grown in serum-free medium for 48-72 hours then replaced every 12 hours to avoid cell cytotoxicity due to the pH change.

#### S phase

cells were arrested at S phase by first arresting cells at G1 phase then replacing the medium with growth medium supplemented with serum and Hydroxyurea (Sigma, UK) at a final concentration of 1mM and incubated overnight.

#### G2/M phase

cells were grown in serum-free medium for 24 hours followed by replacement with fresh growth medium supplemented with 2μg/mL Nocodazole (Sigma, UK). Growth factors in the medium induce cells to progress to G2/M phase while Nocodazole causes cell arrest at G2/M phase since Nocodazole depolarizes the tubulin in microtubules (Figure 1).

### Detection of cell cycle stage

To determine the distribution of cell cycle stages, non-synchronized and synchronized cell cultures were harvested by gentle scraping. Cellular pellets were washed with Phosphate Buffered Saline (PBS) and fixed with ice-cold 70% Ethanol for 48 hours at 4°C. Fixed cells were washed with PBS+2% goat serum, resuspended in 250μL of Propidium Iodide/RNase solution (FxCycle™) (Life technologies, UK) and incubated for an hour at room temperature. Cells were then washed with PBS+2% goat serum and cell cycle analysis was conducted using a BD FACS Calibur (BD Biosciences, UK).

### Flow cytometry

Non-synchronized and synchronized cells were fixed using 70% Ethanol for 48 hours at 4°C then washed with PBS+2% goat serum (Sigma, UK) and resuspended in 1mL of ice-cold PBS. Tubes contained approximately 1×10^5^ cells. Two tubes served as staining controls (blank and isotype control) and three tubes as positive tests. Positive samples were incubated in primary antibodies for 1 hour at 4°C followed by 30 minutes incubation in secondary antibodies. Cells were then incubated in 250μL of Propidium iodide/RNase solution for 1 hour at room temperature prior to flow cytometry analysis. Samples were analyzed using a BD FACS Calibur.

### Immunocytochemistry using Premo™ (FUCCI) Cell cycle Sensor (BacMam 2.0)

A fluorescence ubiquitination cell cycle indicator (FUCCI) was used according to the manufacturer`s instructions (Life Technologies, UK) to assess cell cycle progression. Cell lines were transfected with the BacMam 2.0 gene delivery system which combines two main cell cycle regulators: Cdt1-tagged with red fluorescent protein (RFP) and geminin-tagged with green fluorescent protein (GFP). Cells in G0/G1 phase expressed Cdt1-tagged with RFP were visualised as cells with red nuclei in G0/G1 phase. Cells in S phase co-expressed Cdt1-RFP and geminin-GFP and were visualised with yellow nuclei. Geminin-GFP is predominately expressed in cells during G2/M phase allowing cells with green nuclei to be observed. Briefly, 1×10^3^ BacMam2.0™ particles were diluted in 200μL serum free Opti-MEM™ (Gibco, UK) followed by addition of 1×10^3^ cells and incubated for 10 minutes. Treated cells were seeded on 10mm sterile coverslips in 48-well plates followed by a 48-hour incubation. Cells were fixed with 4% paraformaldehyde (Sigma, UK) and non-specific antigens were blocked with 10% goat serum (Sigma, UK). CD15 and CD15s primary antibodies were applied for one hour followed by incubation in secondary conjugates for 30 minutes. Cells were counterstained with 10mM Hoechst blue (Sigma, UK).

### Confocal microscopy

ICC images were obtained using X40 and X100 (oil immersion) objectives via a Zeiss LSM 510 Meta Axioskop2 confocal microscope using lasers with excitation wavelengths of 405nm (blue), 488nm (green), 568nm (red) and 674 (purple) with diode, argon and HeNe1 lasers respectively. Identical settings were used to image negative controls in which primary antibody was replaced with non-specific Isotype. Semi-quantitative analysis of antigen intensity was measured using Zeiss Zen image software.

### Statistical analysis

All experiments were performed in triplicate and data was expressed as +/− SE. Statistical analyses were performed using one-way ANOVA followed by Tukey’s multiple comparison post-hoc tests using Graph Pad Prism 6 software.

## Author contributions

SAJ, ZM, GJP and HLF designed study and analysed data. SAJ and PC performed experiments. All authors contributed to writing of manuscript. This work was part of SAJ’s PhD thesis.

## Acknowledgements

We would like to thank Mrs Katie F. Loveson who provided technical support for DNA authentication of cell lines used.

## Funding

SAJ was funded by the High Committee of Education Development in Iraq (HCED-IRAQ). ZM was funded by a grant from Animal Free Research UK to GJP. GJP and HLF supported by BTR.

All authors report that there is no conflict of interest.

All cell lines established *in house* were established in accordance with the National Research Ethics Service (NRES) instructions and conformed to the Ethics permission 11/SC/0048.

