## Supplemental data for "CD15 and CD15s expression is associated with G1 phase of the cell cycle in glioma cell lines"

### Supplemental Figure 1

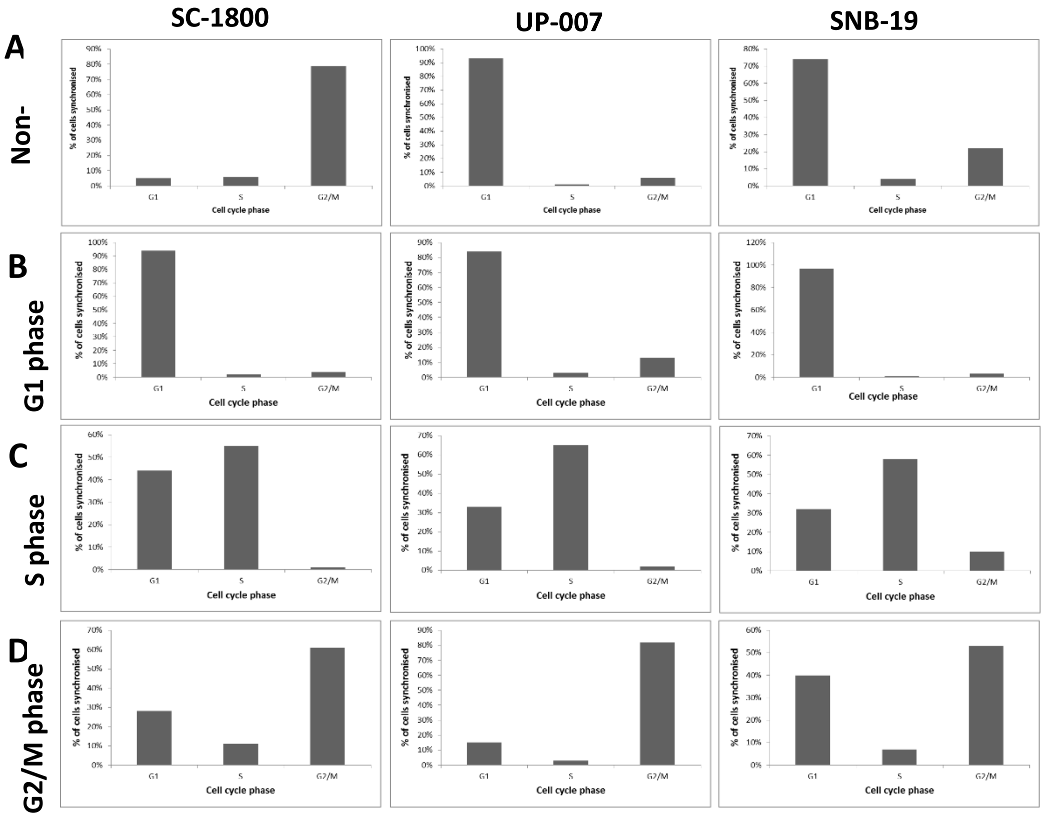

**Supplemental Figure 1:** Synchronisation of cell cultures. Cell cultures were synchronised at different cell cycle stages using serum depletion to arrest cells at G1 phase, 1mM/L hydroxyurea for S phase and 2µg/mL Nocodazole for G2/M phase. Flow cytometry analysis was used to assess the efficiency of cell synchronisation and the distribution of arrested cells in each phase. **(a)** In non-synchronised conditions, the majority of the SC-1800 cells were in G2/M phase while in glioma cell lines, most cells were in G1 phase as determined by flow cytometry analysis of propidium iodide stained cells. **(b)** In serum depletion, cells were arrested at G1 phase at 94% in SC-1800, 84% in UP-007 and 97% in SNB-19. **(c)** Cells incubated with 1mM/mL of Hydroxyurea led to most of cells synchronised in S phase with 55% of SC-1800 cells, 65% of UP-007 and 58% of SNB-19. **(d)** Cells induced to enter G2/M phase then treated with 2µg/mL Nocodazole as a mitotic inhibitor for 24 hours showed that 61% of SC-1800 cells, 82% of UP-007 and 53% of SNB-19 were arrested at G2/M phase.

**Supplemental Figure 2**

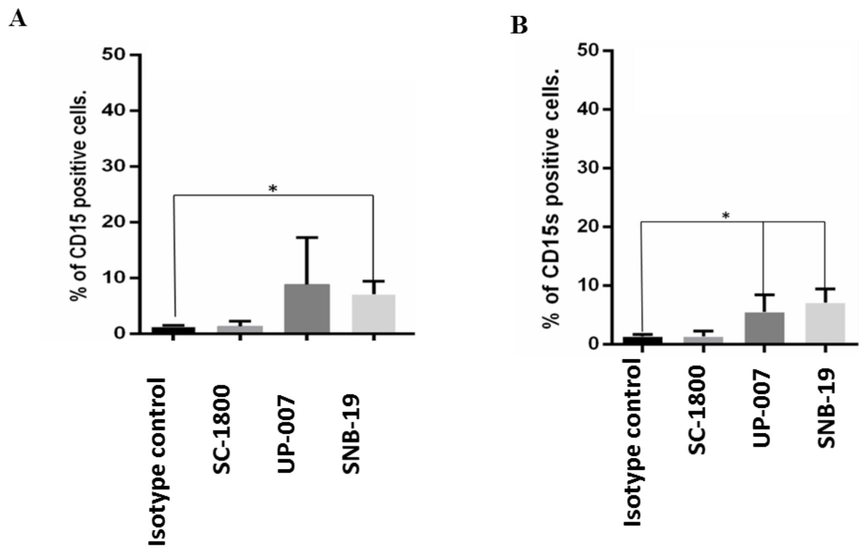


**Supplemental Figure 2:** Expression of CD15 **(a)** and CD15s **(b)** in non-synchronised cells measured using flow cytometry. Quantitative flow cytometric analysis of CD15 **(a)** and CD15s **(b)** expression in SC-1800, UP-007 and SNB-19 with least expression in SC-1800 compared to IgM isotype control *(P≤0.05). Results are representative of three independent experiments carried out in triplicate (n=3).
